# Autotaxin impedes anti-tumor immunity by suppressing chemotaxis and tumor infiltration of CD8^+^ T cells

**DOI:** 10.1101/2020.02.26.966291

**Authors:** Elisa Matas-Rico, Elselien Frijlink, Irene van der Haar Àvila, Apostolos Menegakis, Maaike van Zon, Andrew J. Morris, Jan Koster, Fernando Salgado-Polo, Sander de Kivit, Telma Lança, Antonio Mazzocca, Zoë Johnson, John Haanen, Ton N. Schumacher, Anastassis Perrakis, Inge Verbrugge, Joost van den Berg, Jannie Borst, Wouter H. Moolenaar

**Affiliations:** Division of Biochemistry, Netherlands Cancer Institute, Amsterdam, The Netherlands; Division of Tumor Biology and Immunology, Netherlands Cancer Institute, Amsterdam, The Netherlands; Division of Cell Biology, Netherlands Cancer Institute, Amsterdam, The Netherlands; Division of Molecular Oncology and Immunology, Netherlands Cancer Institute, Amsterdam, The Netherlands; Division of Cardiovascular Medicine, Gill Heart Institute and Lexington Veterans Affairs Medical Center, University of Kentucky, Lexington, KY, US; Amsterdam UMC, Department of Oncogenomics, Amsterdam, The Netherlands; Interdisciplinary Department of Medicine, University of Bari School of Medicine, Bari, Italy; iOnctura SA, Campus Biotech Innovation Park, Geneva, Switzerland

## Abstract

Autotaxin (ATX) is secreted by diverse cell types to produce lysophosphatidic acid (LPA) that regulates multiple biological functions via G protein-coupled receptors LPAR1-6. ATX/LPA promotes tumor cell migration and metastasis mainly via LPAR1; however, its actions in the tumor immune microenvironment remain unclear. Here, we show that ATX secreted by melanoma cells is chemorepulsive for tumor-infiltrating lymphocytes and circulating CD8^+^ T cells *ex vivo*, with ATX functioning as an LPA-producing chaperone. Mechanistically, T-cell repulsion predominantly involves G*α*_12/13_-coupled LPAR6. Upon anti-cancer vaccination of tumor-bearing mice, ATX does not affect the induction of systemic T-cell responses but suppresses tumor infiltration of cytotoxic CD8^+^ T cells and thereby impairs tumor regression. Moreover, single-cell data from patient samples are consistent with intra-tumor ATX acting as a T-cell repellent. These studies highlight an unexpected role for the pro-metastatic ATX-LPAR axis in suppressing CD8^+^ T-cell infiltration to impede anti-tumor immunity, suggesting new therapeutic opportunities.

## Introduction

Efficient infiltration of effector T cells into tumors is associated with positive outcome in several cancer types and determines the response to immunotherapies (Fridman et al., 2017; Ribas and Wolchok, 2018). Chemokines through their G protein-coupled receptors (GPCRs) are major drivers of T-cell migration into tumors, thereby playing a crucial role in the immune response to cancer and influencing tumor fate (Jacquelot et al., 2018; Nagarsheth et al., 2017; Ozga et al., 2021). However, tumors develop various strategies to exclude T cells and suppress T-cell-mediated immunogenicity, for example via tumor-intrinsic silencing of chemokine genes and production of immunosuppressive cytokines (Batlle and Massague, 2019; Joyce and Fearon, 2015; Kerdidani et al., 2019; Spranger et al., 2015; Spranger and Gajewski, 2018). Yet, our understanding of factors that regulate the trafficking of tumor-infiltrating lymphocytes (TILs), either positively or negatively, is incomplete and requires identification of new tractable targets (Anandappa et al., 2020; van der Woude et al., 2017). Here we explore a role for autotaxin (ATX) in this process.

ATX (encoded by *ENPP2*) is a unique lysophospholipase D (lysoPLD) that is secreted by diverse cell types to produce the GPCR agonist lysophosphatidic acid (LPA) from abundantly available lysophosphatidylcholine (LPC) (Perrakis and Moolenaar, 2014; Tokumura et al., 2002; Umezu-Goto et al., 2002). ATX was originally isolated as an “autocrine motility factor” secreted by melanoma cells and characterized as a metastasis-enhancing phosphodiesterase (Nam et al., 2001; Stracke et al., 1992). It is now clear that the ATX-LPA axis plays a key role in a wide variety of (patho)physiological processes, ranging from vascular development (van Meeteren et al., 2006) and lymphocyte homing (Kanda et al., 2008) to tumor progression (Benesch et al., 2018; Mills and Moolenaar, 2003). LPA signals through six GPCRs, termed LPAR1-6 or LPA1-6, showing both distinct and shared signaling activities (Yanagida et al., 2013; Yung et al., 2014). LPAR1-3 belong to the EDG family of GPCRs alongside the sphingosine 1-phosphate (S1P) receptors, whereas the disparate LPAR4-6 members are related to purinergic receptors (Hisano and Hla, 2019; Yanagida et al., 2013). Importantly, ATX binds to cell-surface integrins and heparan sulfate proteoglycans to facilitate delivery of LPA to its cognate receptors in a highly localized manner (Fulkerson et al., 2011; Hausmann et al., 2011; Houben et al., 2013; Kanda et al., 2008).

Numerous studies have documented a critical role for ATX and/or LPA in stimulating cell migration, tumor cell dispersal, invasion and metastasis, mediated primarily by LPAR1 (Auciello et al., 2019; David et al., 2010; Lin et al., 2019; Liu et al., 2009; Marshall et al., 2012). LPAR1 also mediates the recruitment and activation of fibroblasts, a prototypic ATX-secreting cell type, and the ATX-LPAR axis is implicated in promoting tissue fibrosis (Ledein et al., 2020; Sakai et al., 2019; Tager et al., 2008). In addition, activated fibroblasts constitute a large part of solid tumors in which they generate extracellular matrix and contribute to metastasis (Kalluri, 2016; Winkler et al., 2020). It is further important to note that LPAR1-3 commonly mediate enhanced cellular responses, whereas non-EDG receptors LPAR4-6 can exert counter-regulatory actions in that they suppress the migration and invasion of diverse cell types, depending on their dominant G protein-effector pathways (Jongsma et al., 2011; Lee et al., 2008; Takahashi et al., 2017).

ATX has also an important role in the immune system, as it is abundantly expressed in high-endothelial venules (HEVs) that control lymphocyte entry into lymphoid tissue (Kanda et al., 2008; Takeda et al., 2016). Acting through LPAR2, HEV-secreted ATX promotes the random motility of naïve T cells to enhance their transmigration from blood into secondary lymphoid organs and thereby contributes to the control of systemic T-cell responses (Bai et al., 2013; Kanda et al., 2008; Knowlden et al., 2014; Takeda et al., 2016). Thus, the pro-metastatic ATX-LPAR axis regulates the migratory activities of both tumor cells and T cells. However, its actions in the tumor immune microenvironment remain unclear, particularly its dominant LPAR signaling pathways and how ATX/LPA may affect antigen-specific T-cell responses and effector T-cell activity in a tumor context.

Here, we show that ATX/LPA antagonizes the migration of patient-derived TILs and healthy blood-derived CD8^+^ T cells *ex vivo*, and uncover LPAR6 as a T-cell migration inhibitory receptor. By eliciting a robust immune response after anti-cancer vaccination of tumor-bearing mice, we demonstrate that secreted ATX antagonizes tumor infiltration of cytotoxic CD8^+^ T cells and thereby impedes tumor control in a therapeutic setting. Concordantly, in patient samples, single-cell analysis shows an inverse correlation between intratumoral ATX expression and CD8^+^ T cell infiltration. Hence, by revealing ATX/LPA as a suppressor of anti-tumor immunity, our findings shed new light on its multifaceted actions in the tumor microenvironment that could be exploited therapeutically.

## Results

### Through LPA production, ATX secreted by melanoma cells is chemorepulsive for TILs and peripheral CD8^+^ T cells

Melanoma cells are known for their high ATX expression levels among many cancer cell lines and solid tumors (**Figure S1A,B**). This feature is unrelated to genetic changes (www.cbioportal.org), but rather reflects high ATX expression in epidermal melanocytes, a highly motile cell type. We set out to examine how melanoma cell-secreted ATX affects the migration of *ex vivo* expanded melanoma TILs and peripheral blood CD8^+^ T cells. Patient-derived TILs constitute a heterogeneous population of T cells in distinct functional states and other immune cells (Li et al., 2019). During their *ex vivo* expansion driven by anti-CD3 antibody and IL-2 (see Methods) TILs become enriched in CD8^+^ and CD4^+^ T cells and are then used for adoptive TIL therapy in patients (Rohaan et al., 2019).

We first analyzed the effects of LPA and ATX/LPC on the transwell migration of TILs (isolated from two patients). As a positive control, we used chemokine CXCL10 that signals via CXCR3 to promote effector T-cell migration and is implicated in enhancing cancer immunity (Groom and Luster, 2011; Nagarsheth et al., 2017). Strikingly, LPA strongly suppressed the basal migration rate of TlLs (up to 5-fold in patient #1) when assayed over a period of 2 hrs, in a concentration-dependent manner (**Figure 1A,B**). LPA was capable of antagonizing TIL migration towards CXCL10 (**Figure 1C**). LPA was also chemo-repulsive for peripheral blood CD8^+^ T cells isolated from healthy donors (**Figure 1D**). When TILs or CD8^+^ T cells were exposed to recombinant ATX (20 nM) together with its substrate LPC (1-5 *μ*M), their transwell migration was similarly suppressed (**Figure 1E**).

**Figure 1.**
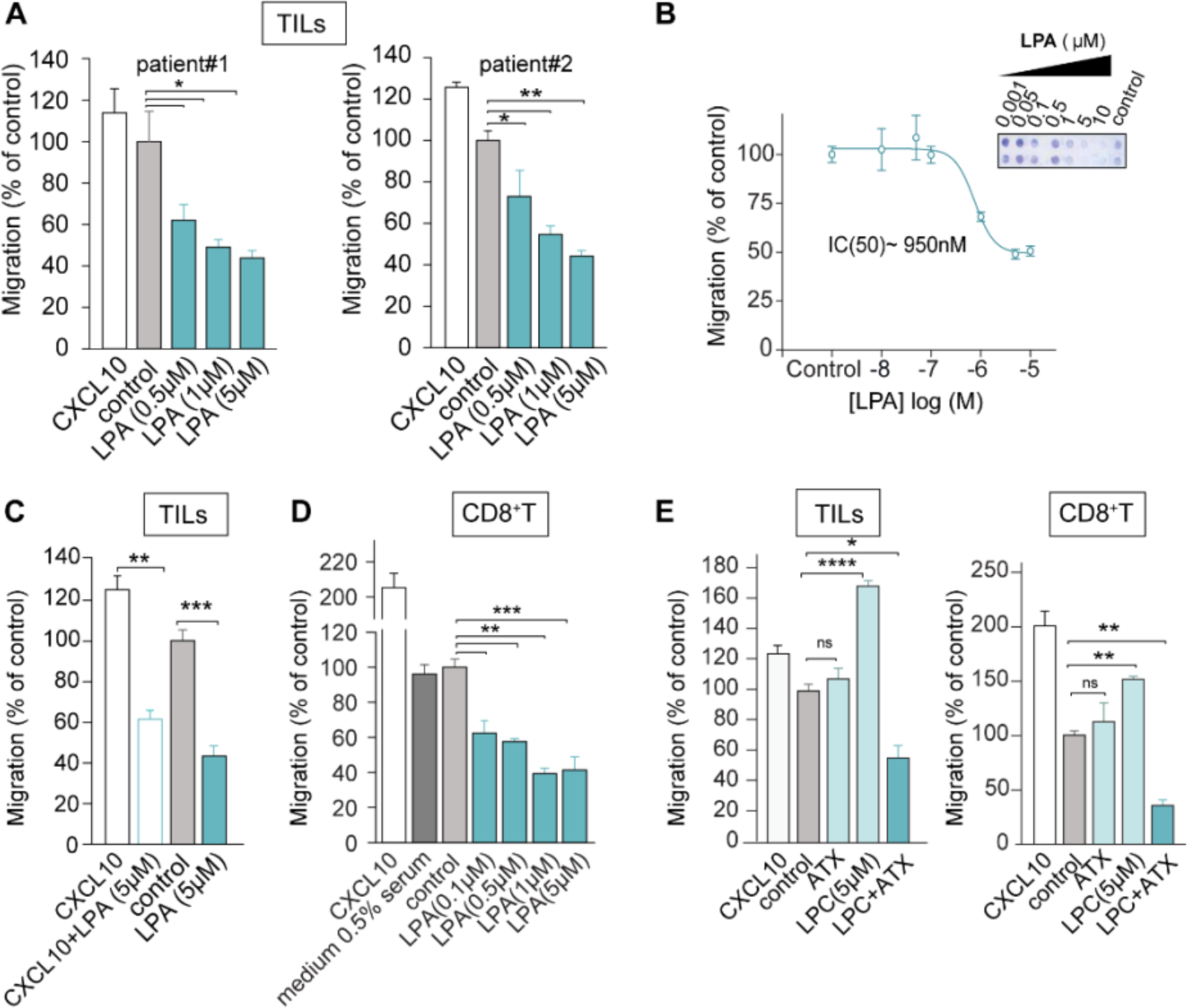
LPA and ATX/LPC are chemorepulsive for TILs and peripheral CD8^+^ T cells. **(A)** Transwell migration of *ex vivo* expanded TILs from two melanoma patients stimulated with LPA(18:1) at the indicated concentrations. Chemokine CXCL10 (1 μM) was used as positive control; “control” refers to serum-free medium. Agonists were added to the bottom wells and incubation was carried out for 2 hrs at 37°C. **(B)** LPA dose-dependency of migration. The inset shows a representative transwell filter after staining. Migration was quantified by color intensity using Image J. **(C)** LPA overrules CXCL10-induced TIL chemotaxis. LPA(18:1) was added together with CXCL10 at the indicated concentrations. **(D)** Migration of CD8^+^ T cells isolated from peripheral blood, measured in the absence (control) and presence of the indicated concentrations of LPA(18:1). Note that the presence of 0.5% serum has no effect. **(E)** Recombinant ATX (20 nM) added together with the indicated concentrations of LPC(18:1) recapitulates the inhibitory effects of LPA(18:1) on TILs and CD8^+^ T cells. **(A-E)** Results are representative of three independent experiments each performed in technical triplicates and expressed as means ± SEM; *p<0.05, **p<0.01, ***p<0.001 (unpaired Student’s t test).

We next analyzed melanoma cell supernatants for their modulatory activity on T cell migration. In addition, we measured concurrently secreted ATX protein and lysoPLD activity. Culture media (containing 0.5% serum) conditioned by melanoma cells (MDA-MB-435 and A375) for 24 hrs markedly suppressed the basal migration and CXCL10-induced chemotaxis of TILs and peripheral CD8^+^ T cells (**Figure 2A**). Secreted ATX protein was readily detected by immunoblotting (**Figure 2B**), while lysoPLD activity was detected simultaneously (**Figure 2C**). By contrast, conditioned media from either ATX knockdown melanoma cells (**Figure 2D**) or ATX-deficient MDA-MB-231 breast carcinoma cells (**Figure 2F**) lacked chemorepulsive activity (**Figure 2E,F**). TIL migration could be rescued by incubating melanoma media with established ATX inhibitors, notably PF-8380 and IOA-289 (formerly CRT750; (Shah et al., 2016)) (**Figure 2G**). Together, these results show that LPA-producing ATX released from melanoma cells is a major T-cell repellent.

**Figure 2.**
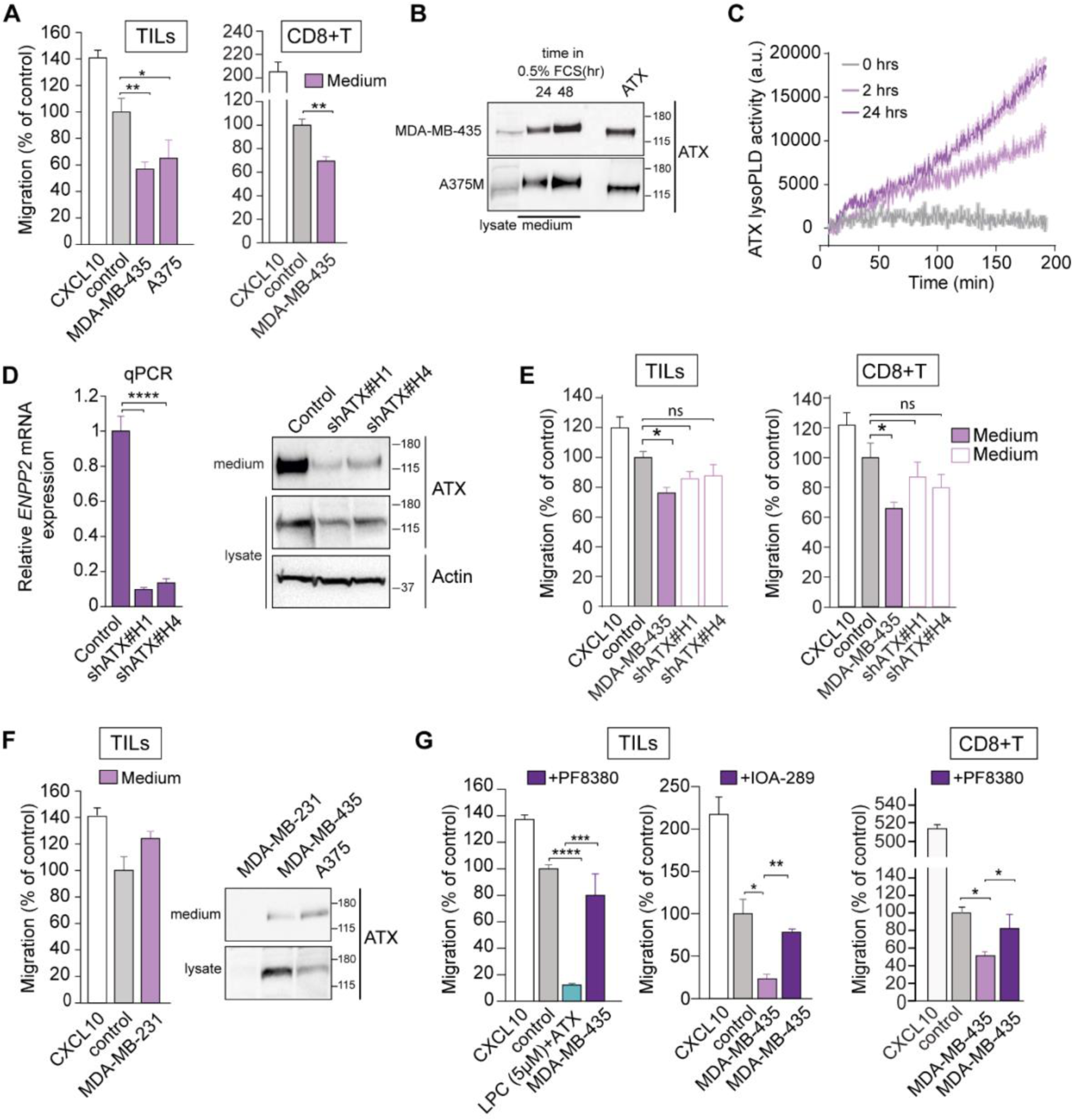
ATX secreted by melanoma cells repels TILs and peripheral CD8^+^ T cells. **(A)** Melanoma-conditioned medium from MDA-MB-435 cells (collected after 24 hrs) is chemorepulsive for TILs and blood-derived CD8^+^ T cells. Experimental conditions as in Figure 1. **(B)** Immunoblot showing ATX expression in medium and cell lysates of MDA-MB-435 and A375M melanoma cells. Cells were incubated in DMEM with 0.5% FCS for 24 or 48 hrs. Recombinant ATX (20 nM) was used as positive control (right lane). **(C)** LysoPLD activity accumulating in melanoma-conditioned media over time. Medium from MDA-MB-435 cells was collected after 2 and 24 hrs, and lysoPLD activity was measured as choline release from added LPC(18:1). **(D)** ATX (*ENPP2*) mRNA expression (relative to cyclophilin) in control and *ENPP2*-depleted MDA-MB-435 cells stably expressing short hairpin (sh)RNAs against ATX. Maximal *ENPP2* knockdown was obtained with shRNA #1 and #4 (of 5 different hairpins). (<u>Right</u>) Immunoblot analysis of ATX expression using shRNA #1 and #4. Actin was used as loading control. **(E)** Melanoma-conditioned medium from ATX knockdown MDA-MD-435 cells (collected after 24 hrs) lacks chemorepulsive activity for CD8^+^ T cells and TILs. **(F)** Conditioned media from ATX-deficient MDA-MB-231 breast carcinoma cells lack chemo-repulsive activity for TILs compared to media from ATX-expressing melanoma cells (MDA-MB-435 and A375; cf **2A**). <u>Right panel</u>, ATX immunoblots from the indicated media and cell lysates. **(G)** ATX inhibition restores the migration TILs and CD8^+^ T cells exposed to melanoma cell-conditioned media. Cells were plated at day 0 in medium containing 10% FCS. After 16 hrs, cells were exposed to medium containing 0.5% FCS and ATX inhibitors (PF-8380 or IOA-289). Conditioned media were collected after 24 hrs. **(A, D-G)** Representative data of three independent experiments each performed in triplicate. Values are expressed as mean ± SEM; *p<0.05 **p<0.01,***p<0.001, ****p<0.0001 (unpaired Student’s t-test).

### ATX as an LPA-producing chaperone

We investigated the relationship between ATX-mediated T-cell repulsion and extracellular LPA levels. It is well established that LPA in freshly isolated plasma increases to high levels due to constitutive ATX-catalyzed LPC hydrolysis (Aoki et al., 2008). The extracellular LPA pool comprises distinct molecular species that differ in their acyl chain composition and binding affinity for individual LPA receptors (Yung et al., 2014). We measured LPA species in media from melanoma cells conditioned at 0, 24 and 48 hrs using LC-MS/MS (Kraemer et al., 2019) (**Figure 3A**). LPA(12:0), (16:0), (18:0), (18:1) and (20:4) were the predominant species in media containing 0.5% serum (**Figure 3B)**. Remarkably, total LPA in TIL-repulsive media declined to very low levels within 24 hrs, despite the fact that ATX activity increased concurrently (**Figures 3C,E** and **2B,C**); by contrast, the corresponding LPC species in these media remained constant or increased slightly over time **(Figure 3D)**. Hence, the loss of LPA in the face of ATX activity is not due to substrate depletion. Rapid depletion of extracellular LPA by melanoma cells has been reported previously (Muinonen-Martin et al., 2014) and is due to its degradation by cell-associated lipid phosphate phosphatases (Sciorra and Morris, 2002). That ATX is fully bioactive at physiologically negligible LPA levels is explained by the fact that ATX binds LPA in its “exit tunnel”, where it is protected from degradation for delivery to its receptors (Keune et al., 2016; Moolenaar and Perrakis, 2011; Nishimasu et al., 2011; Salgado-Polo et al., 2018). These results support the notion that ATX functions as an LPA carrier or LPA-producing “chaperone”.

**Figure 3.**
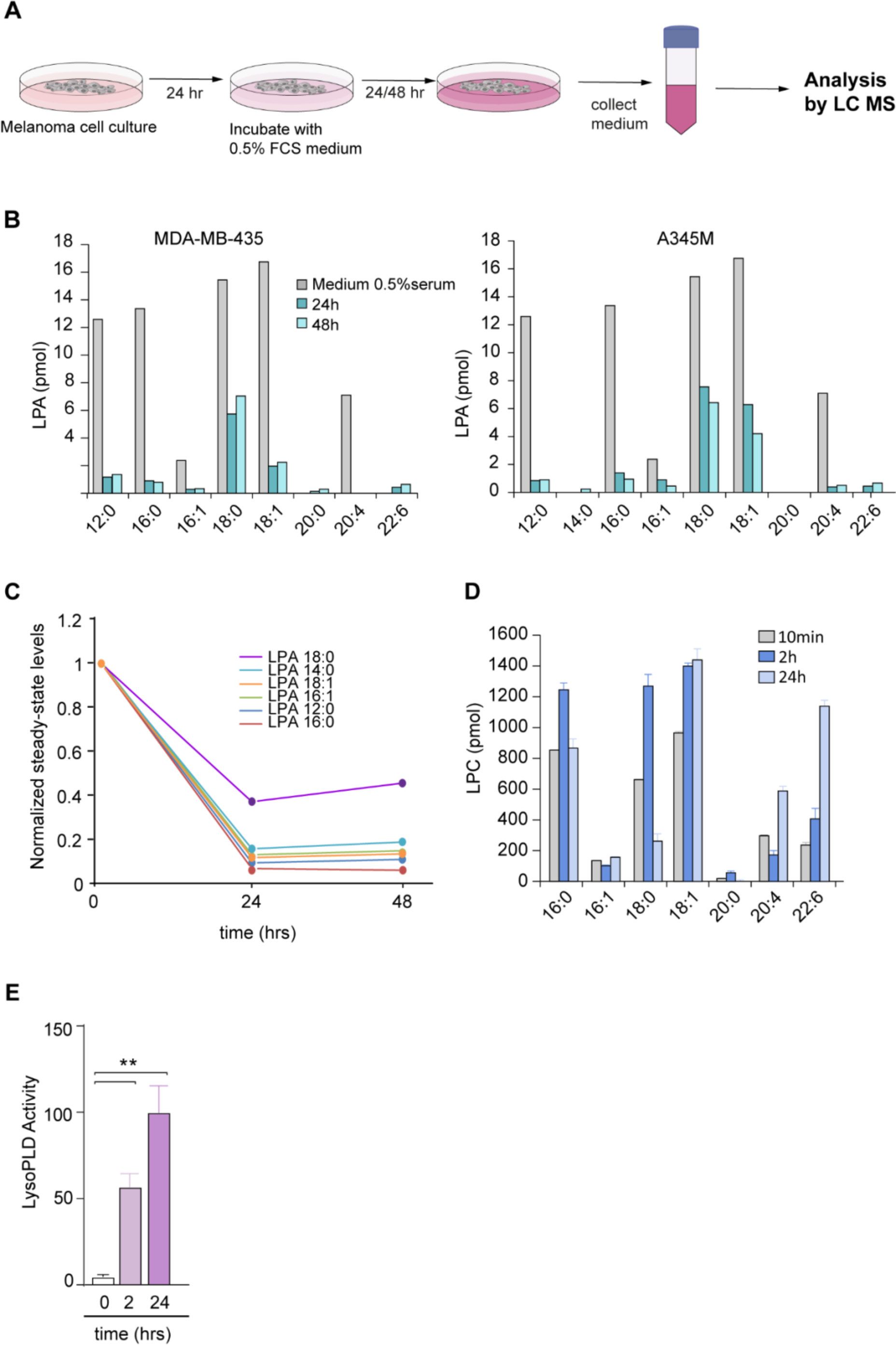
Lysolipid species and secreted lysoPLD activity in conditioned media from melanoma cells. **(A)** Preparation of cell-conditioned media. Melanoma cells in 10-cm dishes were cultured for 24 hrs, washed and then incubated in DMEM containing 0.5% FC. Media were harvested after 24 and 48 hrs, and centrifuged to remove cell debris. LPA species were measured using LC/MS/MS. **(B)** Determination of LPA species in conditioned medium from MDA-MB-435 and A375 melanoma cells, measured at t=0, 24 and 48 hrs, using LC/MS/MS. Predominant serum-borne LPA species are (12:0), (16:0), (18:0), (18:1) and (20:4). Note LPA depletion from the medium (within 24 hrs) upon incubation with ATX-secreting melanoma cells. **(C)** Time-dependent decline of the indicated serum-borne LPA species by melanoma cells. Graph shows normalized steady-state LPA levels in conditioned media from MDA-MB-435 cells. **(D)** LPC species in conditioned medium from MDA-MB-435 cells, measured at t=10 min, 2 hrs and 24 hrs, using LC/MS/MS. Note that LPC levels tend to increase over time. Values from one experiment performed in triplicate and expressed as mean ± SEM. **(E)** Secreted lysoPLD activity increases over time. Medium from MDA-MB-435 cells was collected after 2 and 24 hrs, and lysoPLD activity was measured as choline release from added LPC(18:1). Values from three independent experiments each performed in triplicate and expressed as mean ± SEM; **p<0.01 (unpaired Student’s t-test).

### TIL repulsion involves LPAR6

The T-cell repelling activity of ATX/LPA markedly contrasts to its chemotactic activity for tumor cells, strongly suggesting involvement of different LPA receptors. We examined the LPAR expression repertoire in TILs and blood-derived CD8^+^ T cells using qPCR. *Ex vivo* expanded melanoma TILs (isolated from eight patients) consistently express high levels of *LPAR6* in addition to considerably lower levels of *LPAR2*; an identical pattern was detected in ovarian carcinoma-derived TILs **(Figure 4A**, <u>and data not shown</u>). LPAR6 was also the predominant non-EDG LPA receptor in peripheral blood CD8^+^ T cells alongside LPAR4 and LPAR5 (**Figure 4B**), in agreement with publicly available data (https://www.immgen.org; https://biogps.org). LPAR4 and LPAR5 may have been lost from TILs during tumorigenesis or their *ex vivo* expansion, scenarios that warrant further investigation. Incubating TILs with a novel xanthylene-based LPAR6 antagonist, named XAA (Gnocchi et al., 2020), partially overcame T-cell repulsion by LPA (**Figure 4C**). We therefore conclude that repulsion of TILs and peripheral blood CD8^+^ T cells primarily LPAR6, without excluding possible additional anti-migratory roles for LPAR4 and LPAR5.

**Figure 4.**
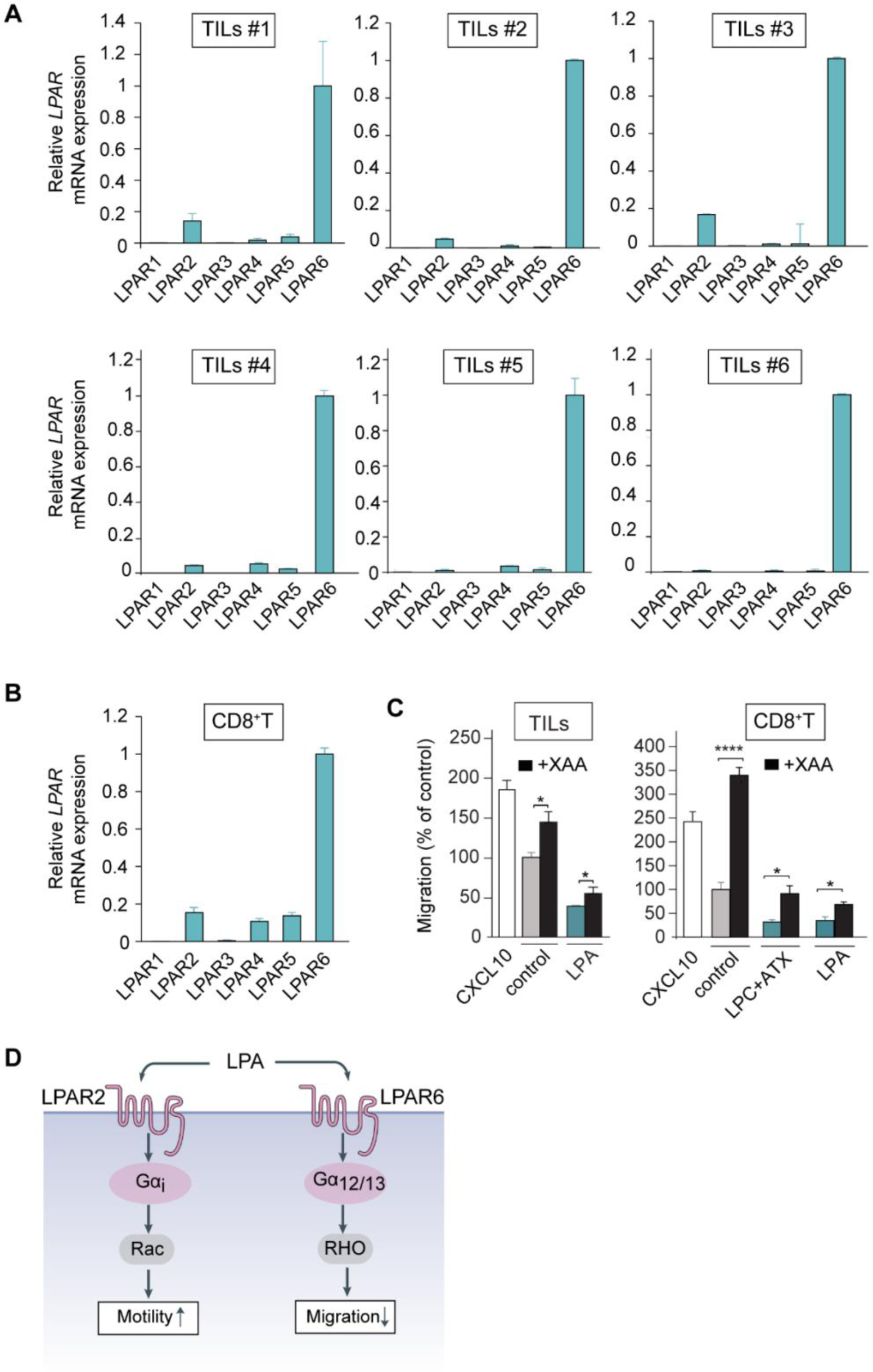
LPAR expression in TILs and peripheral CD8^+^ T cells. **(A)** *LPAR* expression repertoire in *ex vivo* expanded TILs from six patients (qPCR analysis relative to cyclophilin). TIL values are expressed as mean ± SD. **(B)** *LPAR* expression in peripheral CD8^+^ T cells from two healthy donors. Values are expressed as mean ± SD. **(C)** LPAR6 antagonist XAA restores transwell migration of TILs (<u>left panel</u>) and CD8^+^ T cells (<u>right panel</u>) in response to LPA or ATX plus LPC. Conditions as in Figure 1. Cells were treated with XAA (10 μM) or vehicle control (0.5% DMSO) for 24 hrs. Data represent the mean ± SEM of three independent experiments using duplicate samples. *p < 0.05 (unpaired Student’s t-test). **(D)** Schematic illustration of dominant G-protein coupling and signalling outcomes of LPAR2 versus LPAR6.

LPAR6 (P2RY5) preferentially couples to the Gα_12/13_-RhoA pathway that drives cytoskeletal contraction, suppression of cell motility, and other cellular responses (Inoue et al., 2019; Wu et al., 2019; Yanagida et al., 2009; Yung et al., 2014). The function of LPAR6 in T cells has remained largely unexplored despite its high expression in immune cells (https://biogps.org). In contrast to LPAR6, LPAR2 couples to G_i_-mediated Rac GTPase activation and other G protein-effector routes and thereby promotes the random motility of T cells (Kanda et al., 2008; Takeda et al., 2016), as illustrated in **Figure 4D**.

### Impact of ATX on the induction of systemic T-cell responses and tumor infiltration of cytotoxic CD8^+^ T cells

Having shown that ATX through generation and protection of LPA repels TILs and blood-derived CD8^+^ T cells *ex vivo*, we next investigated how ATX affects the anti-tumor T-cell response *in vivo*. We took advantage of an anti-cancer vaccination model using s.c. implanted TC-1 epithelial tumor cells that express the HPV16 E7 oncogene (Lin et al., 1996). TC-1 tumors lack spontaneous T-cell infiltration; however, tumor-specific CD8^+^ T-cell infiltration can be induced by vaccination, as we and others previously described (Ahrends et al., 2016; Ahrends et al., 2017). The DNA vaccine we employed encodes HPV E7 in a gene shuffled configuration to provide a strong MHC class I-restricted CD8^+^ T cell epitope and HPV-unrelated MHC class II-restricted epitopes that elicit CD4^+^ T-cell “help”. These “helped” CD8^+^ T cells have optimal cytotoxic and migratory abilities that allow for effective tumor rejection. Specifically, they readily extravasate and infiltrate into the tumor due to upregulation of chemokine receptors and matrix metalloproteases (Ahrends et al., 2017). This therapeutic setting provides a window to examine the impact of ATX on anti-tumor T-cell responses and tumor rejection.

Since TC-1 cells lack ATX expression, we generated ATX-expressing TC-1 (TC-1^ATX^) cells and confirmed that they secrete active ATX (**Figure S2A, B**). Enforced ATX expression did not significantly alter the growth rate of s.c. injected TC-1 tumor cells (**Figure S2C, D**). This agrees with previous tumor implantation studies showing that ATX-LPAR signaling has little effect on primary tumor growth, but does promote metastasis to distant organs through LPAR1 (David et al., 2010; Lee et al., 2015; Marshall et al., 2012).

#### ATX does not affect induction of systemic T-cell responses

We examined how tumor cell-derived ATX may affect the induction of CD8^+^ and CD4^+^ T-cell responses after vaccination. For this purpose, mice were vaccinated on days 8, 11 and 14 after implantation of wild-type or ATX-expressing TC-1 tumor cells (**Figure 5A**). After vaccination, T cells are primed in the vaccine-draining lymph node from where they egress as differentiated effector T cells into the blood and then infiltrate the tumor via chemotaxis (*e.g.* (Ahrends et al., 2017)). Primed HPV E7-specific CD8^+^ T cells were detected by flow cytometry using H-2D^b^/E7_49-57_ MHC tetramers (Tet) (**Figure 5B**). We monitored vaccine-induced T-cell responses in blood over time (**Figure 5C-E**). The HPV E7-specific systemic CD8^+^ T-cell response measured in blood was similar in TC-1^WT^ and TC-1^ATX^ tumor-bearing mice (**Figure 5C, Figure S3A**), as was the frequency of CD8^+^ T cells with a CD44^+^ CD62L^-^ effector phenotype (**Figure 5D, Figure S3B**). Likewise, the frequency of vaccine-induced CD4^+^ T cells showing a CD44^+^ CD62L^-^ effector phenotype increased to a similar extent in both groups of tumor-bearing mice (**Figure 5E, Figure S3C**). Analysis of the spleens (at day 10 after vaccination) showed no differences in the systemic distribution of HPV E7-specific CD8^+^ T cells (**Figure 5F**), nor in their differentiation into granzyme B (GZB)- and IFN*γ*-expressing cytotoxic T lymphocytes (CTLs) (**Figure 5G**). CD4^+^ T-cell responses in the spleen were also similar between both groups of tumor-bearing mice, as measured by the frequency of IFN*γ*-expressing cells among conventional (FOXP3^-^) CD4^+^ T cells (**Figure S3D,E**). Finally, ATX expression did not influence the frequency of FOXP3^+^ CD4^+^-regulatory T cells (Tregs) (**Figure S3F**). Thus, secreted ATX does not affect the induction of systemic CD8^+^ and CD4^+^ T-cell responses upon vaccination, either in magnitude or quality.

**Figure 5.**
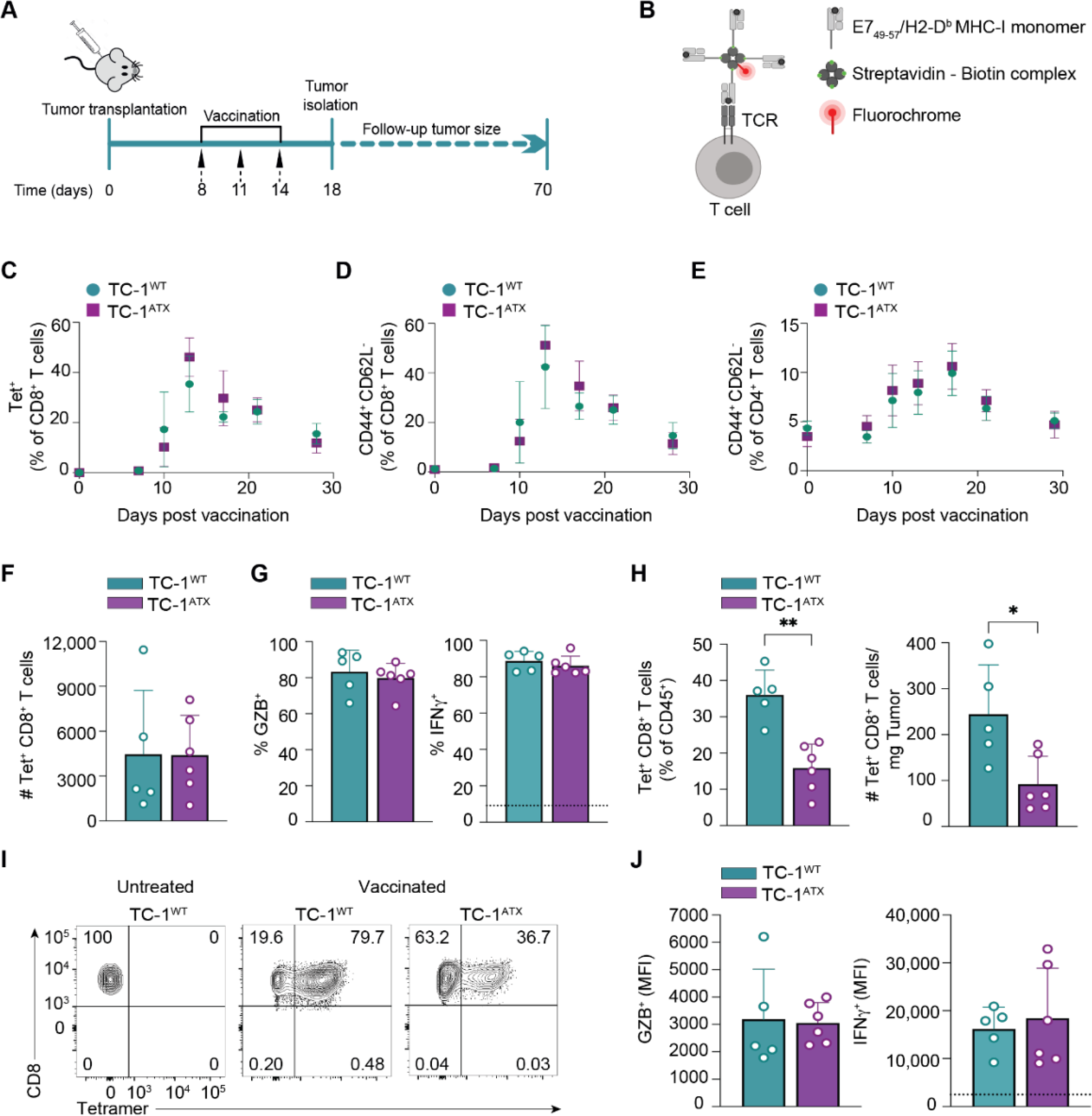
Enforced ATX expression in tumor cells does not affect induction of T-cell responses by vaccination. **(A)** Experimental set-up in the anti-cancer vaccination model. Mice were injected s.c. with wild-type (TC-1^WT^) or ATX-expressing (TC-1^ATX^) tumor cells on day 0, vaccinated on days 8, 11 and 14 and were either sacrificed on day 18, or monitored until day 70. Tumor cells were injected into one flank and the vaccine DNA was “tattooed” into the depilated skin of the opposing flank. Data are from one experiment representative of two experiments. **(B)** The DNA vaccine encodes HPV-E7 protein together with tumor-unrelated helper epitopes. The CD8^+^ T cells that have a TCR specific for the immunodominant E7_49–57_ peptide presented in H-2D^b^ can be detected with MHC class I (MHC-I) tetramers. A tetramer is made by folding E7_49–57_ peptide with MHC-I monomer, conjugating this to biotin and multimerizing it with fluorochrome-conjugated streptavidin. **(C-E)** Monitoring of the T-cell response to vaccination in peripheral blood by flow cytometry in TC-1^WT^ (n=6) and TC-1^ATX^ (n=5) tumor bearing mice. **(C)** Frequency of H-2D^b^/E7_49–57_ tetramer positive (Tet^+^) cells among total CD8^+^ T cells. **(D,E)** Frequency of cells with a CD44^+^CD62L^-^ effector phenotype among total CD8^+^ T cells **(D)** or total CD4^+^ T cells **(E)**. **(F-J)** Analysis of the CD8^+^ T cell response in spleen (**F-J**) and tumor (**H-J**) on day 18 in TC-1^WT^ (n=5) and TC-1^ATX^ (n=6) tumor-bearing mice. **(F)** Absolute number of tetramer positive (Tet^+^) CD8^+^ T cells in spleen. **(G)** Frequency of Granzyme B (GZB)^+^ and IFNᵧ^+^ cells among Tet^+^ CD8^+^ T cells in spleen. IFNᵧ was measured after *ex vivo* PMA/ionomycin stimulation. The dotted line indicates IFNᵧ signal in unstimulated cells. **(H)** Frequency among CD45^+^ hematopoietic cells (left) and absolute number (#, right) of Tet^+^ CD8^+^ T cells in TC-1^WT^ and TC-1^ATX^ tumors. **(I)** Representative flow cytometry plots indicating the percentage of Tet^+^ cells among total CD8^+^ T cells in TC-1^WT^ and TC-1^ATX^ tumors after vaccination and in TC-1^WT^ tumors of non-vaccinated (untreated) mice. **(J)** Mean Fluorescence Intensity (MFI) of GZB^+^ and IFNᵧ^+^ cells within Tet^+^ CD8^+^ T cells in TC-1^WT^ and TC-1^ATX^ tumors. IFNᵧ was measured as in (**G**). **(C-H, J)** Data are expressed as mean ± SD; *p < 0.05, **p < 0.01 (Mann-Whitney U test).

#### ATX repels cytotoxic CD8^+^ T cells from the tumor and impairs tumor control

We then investigated how ATX affects anti-tumor immunity and tumor fate after vaccination. Tumor infiltration of vaccine-induced effector CD8^+^ T cells was analyzed by flow cytometry and immunohistochemistry. Enforced ATX expression significantly reduced the infiltration of HPV E7-specific CD8^+^ T cells into the tumor, in both absolute numbers and frequency among total hematopoietic (CD45^+^) cells (**Figure 5H,I**). ATX did not alter the intrinsic cytotoxicity of the infiltrating CD8^+^ T cells, based on the similar expression levels of GZB and IFN*γ* in Tet^+^ CD8^+^ T cells retrieved from TC-1^WT^ and TC-1^ATX^ tumors (**Figure 5J**). Tumor-derived ATX did not oppose tumor infiltration by conventional (FOXP3^-^) CD4^+^ T cells (**Figure S3G**), nor did it affect their effector quality as inferred from IFN*γ* expression levels (**Figure S3H**). Numbers of infiltrating CD4^+^ Treg cells were also similar between TC-1^WT^ and TC-1^ATX^ tumors (**Figure S3I**). In conclusion, ATX expression by TC-1 tumors impaired infiltration of vaccine antigen-specific CD8^+^ T cells from the blood into the tumor, without affecting CTL quality or infiltration of conventional CD4^+^ T cells and Treg cells into the tumor.

We verified the flow cytometric data by examining T-cell infiltration through quantitative CD8 staining in whole tumor sections by immunohistochemistry. As illustrated in **Figure 6A**, vaccine-induced CD8^+^ T cells were less capable of penetrating ATX-expressing tumors compared to parental tumors. In the parental TC-1^WT^ tumors, CD8^+^ T cells were evenly dispersed throughout the tumor, according to analysis of multiple whole tumor sections. In TC-1^ATX^ tumors, however, CD8^+^ T cells were detected in separate fields, leaving large parts of the tumor non-infiltrated. Quantitative analysis confirmed reduced CD8^+^ T cell infiltration in ATX-expressing tumors (**Figure 6B**). Tumor infiltration of CD4^+^ T cells and Tregs was not affected by ATX expression (**Figure 6C,D**), in agreement with the flow cytometric data.

**Figure 6.**
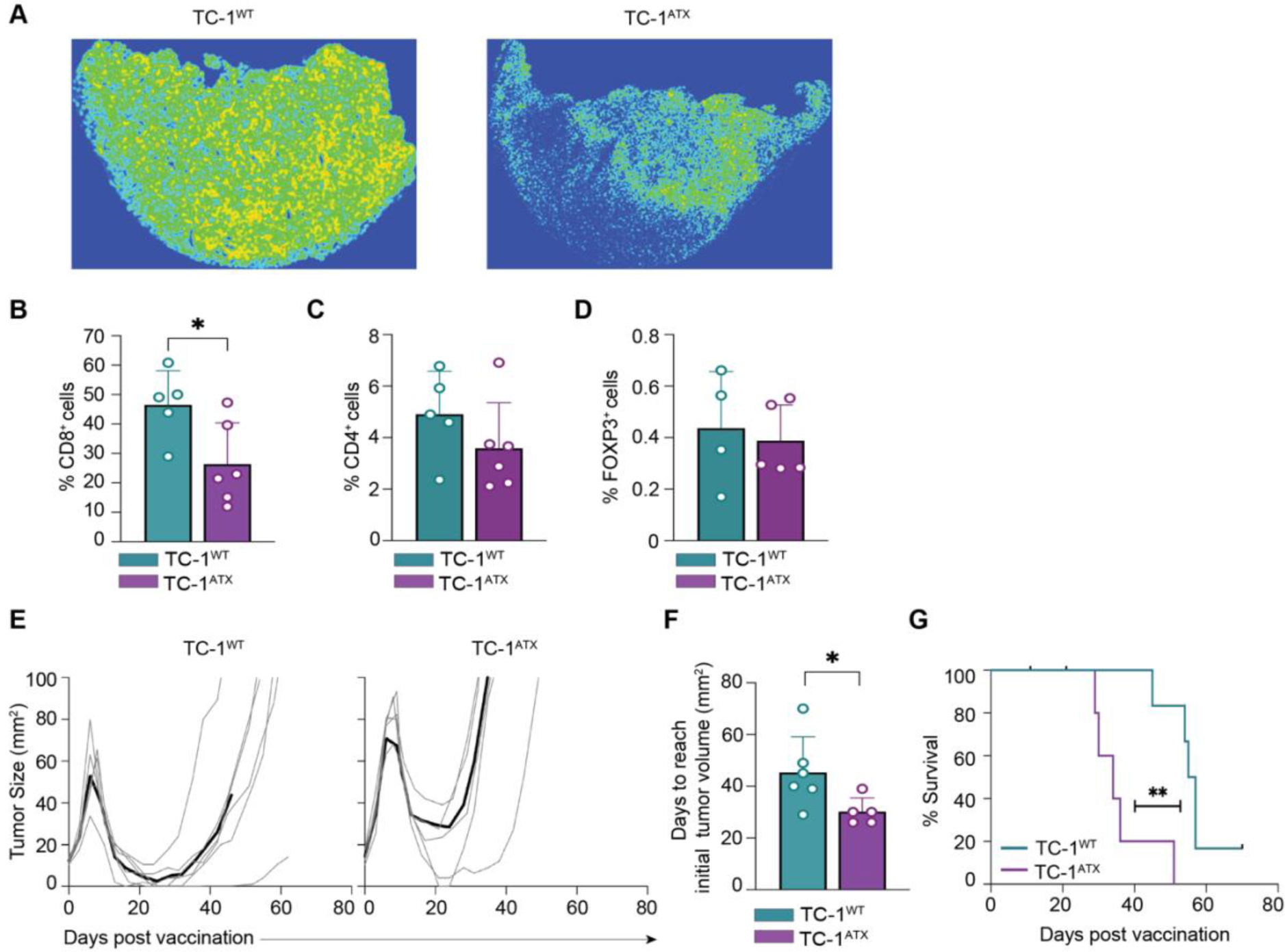
Enforced ATX expression in tumor cells inhibits infiltration of effector CD8^+^ T cells and impedes vaccine-induced tumor control. **(A-F)** Tumor analysis by immunohistochemistry on day 18 in the same mice as analysed in Figure 5. **(A)** Representative heat maps of CD8^+^ immunostainings of tumor sections from vaccinated mice bearing TC-1^WT^ or TC-1^ATX^ tumors. **(B-D)** Quantification in percentages of CD8^+^ **(B,** representative for the data shown in **(A))**, CD4^+^ **(C)** and FOXP3^+^ **(D)** cells out of all nucleated cells as assessed by immunostaining of tumor sections from vaccinated mice bearing TC-1^WT^ or TC-1^ATX^ tumors. (B-D) Data is depicted as mean + SD, *p < 0.05 (Mann-Whitney U test). **(E-G)** TC-1^WT^ (n=6) and TC-1^ATX^ (n=5) tumor-bearing mice received vaccination as outlined in Figure 5 and tumor growth was monitored over time up to day 70. **(E)** Individual growth curves of TC-1^WT^ and TC-1^ATX^ tumors in vaccinated mice. Black lines represent group average. **(F)** Tumor growth delay following vaccination, expressed as number of days required to reach a tumor size corresponding to that at day 7 (see **E**). Data is depicted as mean + SD, *p < 0.05 (Mann-Whitney U test). **(G)** Overall survival curves of tumor-bearing mice. **p < 0.01 (Mantel-Cox analysis). Data in this figure are from one experiment representative of two independent experiments.

We determined the impact of ATX expression on vaccine-induced TC-1 tumor control, following the scheme of **Figure 5A**. Vaccination of mice bearing either TC-1^WT^ or TC-1^ATX^ tumors initially resulted in tumor regression (**Figure 6E**). However, vaccine-induced growth delay of ATX-expressing tumors was significantly reduced in comparison to TC-1^WT^ tumors, as was the overall survival rate of mice bearing TC-1^ATX^ tumors (**Figure 6F,G**). Collectively, these findings demonstrate that ATX released by tumor cells impairs cytotoxic CD8^+^ T-cell infiltration and dispersion throughout the tumor, and thereby impairs tumor control in a therapeutic setting.

### Intratumoral *ENPP2* expression in melanoma inversely correlates with CD8^+^ T-cell infiltration

Finally, through single-cell RNA-seq analysis, we sought clinical evidence for intratumoral ATX functioning as a CD8^+^ T-cell repellent in human melanoma. Remarkably, abundant *ENPP2* expression is detected not only in melanoma but in virtually all solid tumors (www.cbioportal.org), showing little correlation with *ENNP2* expression in the corresponding cancer cell lines (**Figure S1A,B**). This supports the view that a substantial part of the tumor *ENPP2* transcripts is derived from non-malignant stromal cells, notably cancer-associated fibroblasts (CAFs) and adipocytes known for their high ATX expression, depending on the cancer type (Auciello et al., 2019; Brindley et al., 2020).

We analyzed *ENPP2* expression patterns and CD8^+^ T-cell infiltration using single-cell RNA-seq (scRNA-seq) results from 32 melanoma tumors prior to immunotherapy, in which diverse cell subsets could be distinguished (Jerby-Arnon et al., 2018). *ENPP2* expression in individual cells (n=7186) and its association with CD8^+^ T-cell infiltration was examined in all subsets, namely malignant cells, CD8^+^ and CD4^+^ T cells, B cells, NK cells, CAFs, tumor-associated macrophages and endothelial cells. **Figure 7A** shows the melanoma samples grouped by individual cell types. Whereas lymphocytes do not express ATX, significant *ENPP2* expression was detected not only in malignant cells and CAFs but, remarkably, also in macrophages and endothelial cells (**Figure 7B**). Tumors with the highest intratumoral *ENPP2* expression – in both cancer and stromal cells - contained significantly fewer CD8^+^ T cells, while low *ENPP2* expression was associated with enhanced CD8^+^ T-cell infiltration, as quantified by Pearson’s correlation analysis (r=0.4; p=0.01) (**Figure 7C**).

**Figure 7.**
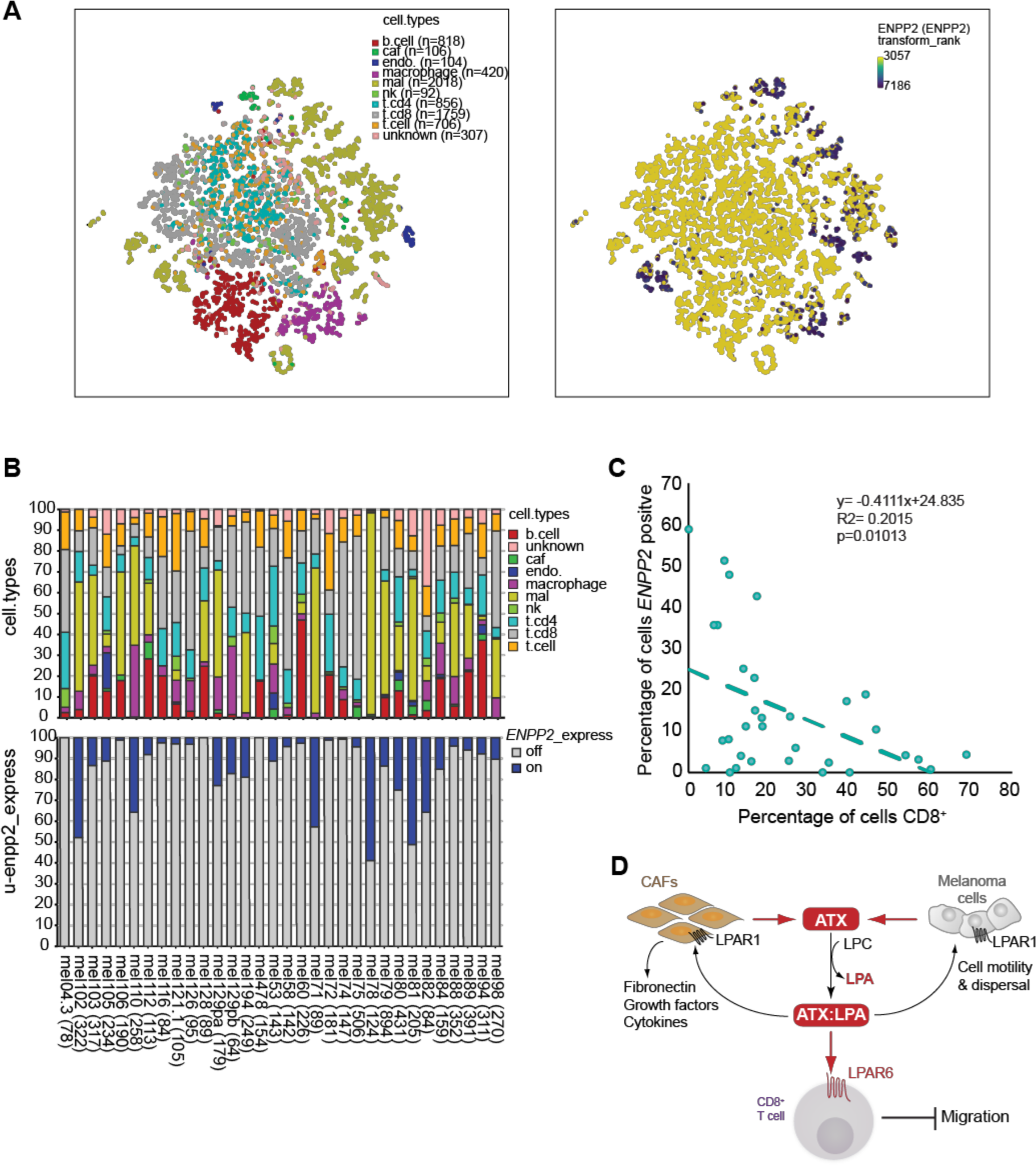
Single-cell analysis of *ENNP2* expression in melanoma tumors and its inverse correlation with CD8^+^ T-cell accumulation. **(A)** tSNE embedding of 7186 single cells (complexity = 5) from 32 melanoma patients as described (Jerby-Arnon et al., 2018). Data were used to project patients, inferred cell types and log_2_ *ENPP2* expression values, respectively, as described in Methods. <u>Right panel</u> shows *ENPP2* expression (blue/purple dots high expression) as overlay on single cells presented in the <u>left panel</u>. Intratumoral *ENPP2* expression is detected in malignant cells (<u>ma</u>l), cancer-associated fibroblasts (<u>ca</u>f), macrophages and endothelial cells (<u>endo</u>), but not in lymphocytes (T, B and NK cells). **(B)** Stacked bar graph showing the percentages of inferred cell type per individual patient sample (top), and the percentage of *ENPP2*-expressing cell types (bottom). **(C)** Negative correlation of intratumoral *ENPP2* expression and CD8^+^ T-cell accumulation. Pearson correlation between the percentage of inferred *ENPP2*-expressing cells and CD8^+^- positive cells (R=0.4; p=0.01). **(D)** Model of the melanoma immune microenvironment. In this model, ATX is secreted by melanoma cells and diverse stromal cells, particularly fibroblasts (CAFs), to convert extracellular LPC into LPA. ATX functions as an LPA-producing chaperone (ATX:LPA) that carries LPA to its GPCRs and exerts dual actions: it suppresses T-cell infiltration through G_12/13_-coupled LPAR6, while it promotes melanoma cell dispersal and activates CAFs via LPAR1 (mainly via G_I_). Activated CAFs release growth factors and produce extracellular matrix. Random T-cell motility mediated by LPAR2 is not illustrated (see Figure 4D). See text for further details.

Elevated *ENPP2* expression in melanoma samples was also associated with reduced CD4^+^ T-cell infiltration, but not with macrophage accumulation **(Figure S4A,B).** ATX-mediated repulsion of CD4^+^ T cells was not observed in the mouse vaccination model; differences in the functional state of the respective CD4^+^ T-cell populations or species-specific receptor expressions might account for this discrepancy. As a caveat, scRNA-seq analyses do not detect all transcripts in every single cell, while *ENPP2* expression is not necessarily predictive of secreted ATX activity. Nonetheless, the single-cell transcriptomics analysis is consistent with our *in vivo* findings, namely that intratumoral ATX repels CD8^+^ T cells from the tumor. Altogether, our findings support a model of intra-tumor ATX/LPA signaling (**Figure 7D**) to be discussed below.

## Discussion

The cellular and molecular mechanisms that contribute to the exclusion of cytotoxic CD8^+^ T cells from the tumor are not well understood, hampering progress in improving immunotherapy efficacy (Anandappa et al., 2020; Joyce and Fearon, 2015; van der Woude et al., 2017). Tumor-intrinsic mechanisms underlying T-cell exclusion involve, for example, transcriptional chemokine silencing (Spranger et al., 2015; Spranger and Gajewski, 2018) and production of immunosuppressive cytokines such as TGFβ (Batlle and Massague, 2019; Mariathasan et al., 2018). However, secreted factors and T-cell GPCRs that counteract TIL infiltration remain to be identified. In this study, we demonstrate that LPA-producing ATX secreted by tumor cells is a major repellent for human TILs and healthy CD8^+^ T cells under *ex vivo* conditions, with a dominant role for the anti-migratory Gα_12/13_-coupled receptor LPAR6. Moreover, we show that secreted ATX repels cytotoxic CD8^+^ T cells from s.c. engrafted tumors to impede anti-tumor immunity and tumor regression in a therapeutic setting.

ATX/LPA is widely known for its chemotactic activities towards both normal and tumor cells, acting via LPAR1, and to enhance the random motility of T cells via LPAR2 (Kanda et al., 2008; Knowlden et al., 2014). Unexpectedly, we initially observed T-cell chemo-repulsive effects of exogenous ATX/LPA and melanoma cell-secreted ATX of TILs and peripheral blood CD8^+^ T cells *ex vivo*, with ATX/LPA antagonizing the migration towards chemokine CXCL10. Whereas CD8^+^ T cells express multiple LPA receptors, the unique LPAR expression pattern in *ex vivo* expanded TILs allowed us to define LPAR6 as the predominant T-cell anti-migratory receptor by default. In this respect, it should be emphasized that biological outcome is determined by the balance in expression of GPCRs that signal mainly via G_i_ (i.e. chemokine and chemotactic EDG receptors LPAR1-3) versus those that couple predominantly to the Gα_12/13_-RhoA pathway, notably anti-chemotactic non-EDG receptors LPAR4-6, as exemplified by the present findings.

Contrary to prevailing notions, tumor cell-secreted ATX failed to raise extracellular LPA levels as its lysoPLD activity was outperformed by cell-associated LPA-degrading activity (**Figure 3**). By binding LPA in its tunnel, ATX can protect LPA from rapid degradation (Keune et al., 2016; Moolenaar and Perrakis, 2011; Nishimasu et al., 2011; Salgado-Polo et al., 2018) and, as such, functions as the physiologically relevant LPA carrier or “chaperone”. Based on its calculated lifetime, the ATX/LPA complex can diffuse over a relatively long distance in the extracellular milieu (Salgado-Polo et al., 2018; Saunders et al., 2011). Precisely how ATX delivers bound LPA to its cell-surface receptors awaits further functional and structural studies.

LPAR6 now joins the few select GPCRs that counteract T-cell chemotaxis through the Gα_12/13_-RhoA pathway. Among them, lysolipid receptor S1PR2 is arguably the best characterized member (Baeyens et al., 2015; Laidlaw et al., 2019), although a role in immuno-oncology has not been documented thus far. LPAR6 (P2RY5) is of special interest as it displays its highest expression in immune cells (https://biogps.org) and is strongly induced upon activation of chicken T cells through as-yet-unknown mechanisms (Kaplan et al., 1993; Webb et al., 1996). Furthermore, LPAR6 prefers 2-acyl-rather than 1-acyl-LPA species as ligand (Yung et al., 2014), which may explain the relatively high IC_50_ value for 1-oleyl-LPA observed in T-cell migration assays (**Figure 1B)**. Although its non-EDG relatives LPAR4 and LPAR5 were not detected in *ex vivo*expanded TILs (**Figure 4A**), the latter receptor is nonetheless of immuno-oncological importance since its genetic deletion in mice enhances T-cell receptor activity and anti-tumor responses (Mathew et al., 2019). To what extent LPAR6 and LPAR5 may act redundantly or synergistically in T-cell signal transmission remains to be investigated.

Building on our *in vitro* findings, we pursued the impact of tumor-intrinsic ATX on T-cell responses in the mouse TC-1 tumor model that is often used in anti-cancer vaccination studies. For this purpose, we stably expressed ATX in TC-1 cells that lack endogenous *Enpp2* expression and confirmed their LPA-producing activity. Vaccination induces the simultaneous activation of CD4^+^ and CD8^+^ T cells to optimize the cytotoxic T-cell response in magnitude and quality (Ahrends et al., 2017). “Helped” CD8^+^ T cells acquire chemokine receptors to increase their migration capacity and enhanced metalloprotease activity that enables them to invade tumor tissue to promote tumor regression (Ahrends et al., 2017; Borst et al., 2018). We established that tumor-intrinsic ATX has no effect on vaccine-induced CD4^+^ and CD8^+^ T-cell responses. The cytotoxic CD8^+^ T cells thus displayed optimal effector capacity independent of ATX activity. Importantly, despite the robust anti-tumor immune response, tumor-intrinsic ATX was capable of significantly impeding tumor infiltration of cytotoxic CD8^+^ T cells and suppressing tumor rejection. These findings highlight a key role for LPA-producing ATX in suppressing anti-tumor immunity in a therapeutic setting. By inference, LPAR6 most likely plays a dominant role in mediating ATX-induced T cell repulsion *in vivo*, possibly in concert with LPAR5, but this needs further investigation. Further development of specific LPAR6 antagonists (Gnocchi et al., 2021) would enable a robust pharmacological characterization and help dissect the ATX-LPAR immune signaling network in further detail.

In a clinical setting, single-cell analysis of melanoma tumors (Jerby-Arnon et al., 2018) showed significant *ENPP2* expression in malignant cells, CAFs, tumor-associated macrophages and endothelial cells, which further accentuates the complexity of ATX/LPA signaling in the tumor microenvironment (**Figure 7A-C**). Consistent with our *in vivo* findings, high intratumoral *ENPP2* expression was associated with reduced CD8+ T-cell infiltration. Additionally, *ENPP2* expression inversely correlated with CD4^+^ T-cell accumulation, but not with macrophage recruitment (**Figure S4**), findings that should be validated in future immuno-histochemical analyses of select patient samples.

Taken together with previous evidence, our findings support a simplified model of the tumor (melanoma) microenvironment illustrated in **Figure 7D**. In this model, LPA-producing ATX is secreted by both tumor and stromal cells and - complexed with LPA - counteracts tumor infiltration of CD8^+^ T cells mainly via G_12/13_-coupled LPAR6, while it activates tumor cells and pro-tumorigenic fibroblasts (CAFs) in autocrine/paracrine loops via LPAR1, which signals predominantly via G_i_. ATX/LPA-stimulated tumor cells acquire a pro-metastatic phenotype, while activated fibroblasts drive immune escape by posing a physical barrier to TILs and secreting immunosuppressive molecules (De Jaeghere et al., 2019). As ATX is abundantly expressed in most solid tumors (**Figure S1B**), this model extends beyond melanoma to many cancer types. Since the tumor microenvironment is heterogeneous and cancer type-specific, ATX/LPA signaling outcome will critically depend on the composition and LPAR expression repertoire of the immune cell infiltrate, and likely also on the spatial arrangement of ATX-secreting stromal cells within the tumor.

In conclusion, by suppressing anti-tumor immunity while promoting metastasis via different LPA receptors, the ATX-LPAR signaling axis creates an T-cell excluding, pro-tumorigenic microenvironment that is amenable to therapeutic intervention. Our findings pave the way for addressing major outstanding questions on ATX/LPA in other immunotherapeutic settings, such as genetically engineered melanoma models and/or patient-derived xenografts (PDX) engrafted in humanized mouse models (Patton et al., 2021; Rosato et al., 2018). Such clinically relevant models will provide further insight into the dual pro-tumor actions of ATX; furthermore, they will offer an opportunity to evaluate the anti-tumor benefits of pharmacological ATX inhibition, for example in combination with immune checkpoint inhibitors.

## Methods

### Cell culture and materials

MDA-MB-435 and A375M melanoma cells were grown in Dulbecco’s modified Eagle’s medium (DMEM) supplemented with 10% fetal bovine serum (FBS) at 37°C under 5% CO_2_. Patient-derived TILs cells were grown in Roswell Park Memorial Institute (RPMI) 1640 medium supplemented with 10% human serum at 37°C under 5% CO_2_. Human CD8^+^ T cells were isolated from buffy coats, activated with anti-CD3 and CD28 mAbs that were plate-bound and expanded in RPMI 1640 medium supplemented with 10% human serum and 100 IU/mL IL-2 and 5 ng/ml IL-15 at 37°C under 5% CO2. Interleukins and CXCL10 were from PeproTech. LPA(1-oleoyl) was obtained from Avanti Polar Lipids. Human ATX was produced in HEK293 Flip-in cells and purified as previously described (Salgado-Polo et al., 2018). Fibronectin and PF-8380 were purchased from Sigma-Aldrich. IOA-289 (formerly CRT750) was synthesized as previously described (Shah et al., 2016).

### Isolation and expansion of melanoma-derived TILs

TIL isolation and expansion was started by generation of a single cell suspension by enzymatic digestion of the resected metastatic tumor material obtained by surgery. Resulting cell suspensions were cultured in the presence of 6000 IU/ml IL-2 (Proleukin, Novartis) for two to four weeks. During the subsequent Rapid Expansion Protocol of two weeks, T cells were cultured in 50% RPMI/50% AIM-V medium in the presence of 3,000 IU/ml IL-2, 30 ng/ml anti-CD3 antibody (OKT-3, Miltenyi) and irradiated autologous PBMCs (feeder cells in 200-fold excess over TIL).

### Isolation of peripheral CD8^+^ T cells

Human peripheral blood mononuclear cells (PBMCs) were isolated from fresh buffy coats using Ficoll-Paque Plus (GE Healthcare) gradient centrifugation. Total CD8^+^ T cells were isolated using magnetic sorting with CD8 microbeads (Miltenyi Biotec). Blood samples were obtained from anonymized healthy male donors with written informed consent in accordance to guidelines established by the Sanquin Medical Ethical Committee.

### Conditioned media

Conditioned media were collected from MDA-MB435 and A375M cells. Sub-confluent 10-cm dishes of melanoma cells were washed with PBS and incubated in DMEM containing 0.5% FCS. Conditioned medium was harvested after 24 and 48 hrs, and centrifuged for 30 min at 4000 rpm in a tabletop centrifuge to remove cell debris.

### Transwell migration assays

T cell migration was measured using 48-well chemotaxis chambers (Neuro Probe, Inc.) equipped with 5 µm-pore polycarbonate membranes (8 µm-pore for melanoma cells), which were coated with fibronectin (1 µg/ml). Cells (1×10^6^/ml) were added to the upper chamber. Migration was assessed after 2 hrs for TILs and CD8^+^ T cells at 37°C in humidified air containing 5% CO_2_. Migrated cells were fixed in Diff-Quik Fix and stained using Diff-Quik II. Migration was quantified by color intensity measurements using Image J software.

### ATX lysoPLD activity

ATX enzymatic activity in conditioned media was measured by steady-state choline release from exogenously added LPC using a coupled reaction, as detailed elsewhere (Salgado-Polo et al., 2018). Briefly, media were centrifuged for 45 min at 4,500 rpm, upon which 75 μl of the supernatants were plated on 96-well plates together with 600 μM LPC(18:1), 1 U ml^-1^ choline oxidase, 2 U ml^-1^ horseradish peroxidase (HRP) and 2 mM homovanillic acid (HVA), reaching a final volume of 100 µl. ATX activity was measured by HVA fluorescence at λ_ex_/λ_em_ = 320/460 nm every 30 s for at least 160 min at 37°C with a Pherastar plate reader (BMG Labtech). Since ATX activity *in vitro* presents a ∼15-min lag phase, the subsequent linear slope (60-160 min) was used to perform all analyses. Triplicate measures were statistically analyzed by an unpaired t test.

### Western blotting

Cells were washed in ice-cold PBS (phosphate-buffered saline containing 2 mM Ca^2+^ and Mg^2+^), lysed in RIPA buffer with protease inhibitors and spun down. Equal amounts of proteins were determined by BCA protein assay kit (Pierce), separated by SDS-PAGE using pre-cast gradient gels (4-12% Nu-Page Bis-Tris, Invitrogen) and transferred to nitrocellulose membranes. The membrane was blocked for 1 hr at room-temperature in 5% skimmed milk in TBST. Incubation with antibodies was done overnight at 4°C, followed by 1 hr incubation with horseradish peroxidase-conjugated secondary antibodies (DAKO, Glostrup, Denmark). Proteins were visualized using ECL Western blot reagent (GE Healthcare, Chalfont St. Giles, UK).

### qPCR analysis

Expression levels of LPA receptors and ATX/ENPP2 were quantified by RT-qPCR. Total RNA was isolated using GeneJET purification kit (Fermentas). cDNA was synthesized by reverse transcription from 2 mg RNA with oligodT 15 primers and SSII RT enzyme (Invitrogen). qPCR was performed on a 7500 Fast System (Applied Biosystems) as follows: 95°C for 2 min followed by 40 cycles at 95°C for 15 s followed by 60°C for 1 min. 200 nM forward and reverse primers,16 ml SYBR Green Supermix (Applied Biosystems) and diluted cDNA were used in the final reaction mixture. Cyclophilin was used as reference gene and milliQ was used as negative control. Normalized expression was calculated following the equation NE = 2(Ct target-Ct reference). Primers used: LPA1 forward AATCGGGATACCATGATGAGT, reverse CCAGGAGTCCAGCAGATGATAAA; LPA2 forward CGCTCAGCCTGGTCAAAGACT, reverse TTGCAGGACTCACAGCCTAAAC; LPA3 forward AGGACACCCATGAAGCTAATGAA, reverse GCCGTCGAGGAGCAGAAC; LPA4 forward CCTAGTCCTCAGTGGCGGTATT, reverse CCTTCAAAGCAGGTGGTGGTT; LPA5 forward CCAGCGACCTGCTCTTCAC, reverse CCAGTGGTGCAGTGCGTAGT; LPA6 forward AAACTGGTCTGTCAGGAGAAGT, reverse CAGGCAGCAGATTCATTGTCA; ENPP2 forward ATTACAGCCACCAAGCAAGG, reverse TCCCTCAGAGGATTTGTCAT; Cyclophilin forward CATCTGCACTGCCAAGACTGA and reverse TTGCCAAACACCACATGCTT.

### ATX knockdown melanoma cells

To generate ATX knockdown melanoma cells, we used five human ENPP2 shRNAs in the lentiviral vector pLKO1: (TRC human shRNA library; TRCN0000048993, TRCN0000048995-TRCN0000048997 and TRCN0000174091). To generate particles for stable infections, HEK293T cells were transfected with single shRNA hairpins using the calcium phosphate protocol; the virus particles were collected at 48 hrs after transfection. ENPP2 stable knockdown cells were selected and maintained in medium containing 2 μg/ml puromycin.

### Lipid extraction and LC-MS/MS measurements of LPA

Extraction of lipids from cell-free conditioned media was done using acidified organic solvents and measurement of seventeen LPA species was accomplished using LC-MS/MS. Quantitation of LPA species was achieved using LPA(17:0) as an internal standard. Experimental details can be found elsewhere (Kraemer et al., 2019).

### Studies in mice

Six to eight-week old female C57BL/6JRj (B6) mice were obtained from Janvier Laboratories (Le Genest Saint Isle, France) and maintained in individually ventilated cages (Innovive, San Diego, CA) under specific pathogen-free conditions. All mouse experiments were performed in accordance with institutional and national guidelines and were approved by the Committee for Animal Experimentation at the NKI.

### Tumor cells and subcutaneous transplantation

TC-1 tumor cells are lung epithelial cells engineered to express HPV16 E6 and E7 proteins (Lin et al., 1996). Cells were obtained from the Leiden University Medical Center in 2015, and the authors did not perform further authentication. TC-1 cells stably overexpressing ATX were generated by retroviral transduction and ATX overexpression was validated by Western blotting (Supplementary Figure 3A). TC-1 cells were cultured in medium, supplemented with 10% fetal calf serum (FCS), 0.1 mM non-essential amino acids, 1 mM sodium pyruvate, 2 mM L-glutamine, 10 mM HEPES and antibiotics at 37°C, 5% CO_2_. TC-1 cell stock was tested negative for *Mycoplasma* by PCR, and cells thawed from this stock were used within 3 passages for *in vitro* and *in vivo* experiments. On day 0, mice were anesthetized with isofluorane and injected s.c. with 1× 10^5^ TC-1 tumor cells. Tumor size was measured by calipers in two dimensions and calculated as: area (mm^2^) = width x length. Mice were monitored twice per week. Mice were sacrificed when the tumor diameter reached 15 mm or when the tumor size reached 100 mm^2^. In the survival curves, censored events indicate mice were sacrificed before treated tumors reached 100 mm^2^.

### Anti-cancer vaccination

The HELP-E7SH plasmid DNA vaccine was described previously and validated in detail. Intra-epidermal DNA “tattoo” vaccination was performed as described in the same papers. Briefly, the hair on a hind leg was removed using depilating cream (Veet, Reckitt Benckiser) prior to vaccination, mice were anesthetized and 15 μl of 2 mg/ml plasmid DNA solution in 10 mM Tris, 1 m M EDTA, pH 8.0 was applied to the hairless skin with a Permanent Make Up tattoo device (MT Derm GmbH, Berlin, Germany), using a sterile disposable 9-needle bar with a needle depth of 1 mm and oscillating at a frequency of 100 Hz for 45 sec.

### Tissue preparation and flow cytometry

At the indicated days, blood was sampled from live mice or mice were sacrificed and lymphoid tissues and tumors were harvested. Peripheral blood cells were collected by tail bleeding in Microvette CB300 LH tubes (Sarstedt). Red blood cells were lysed in 0.14 M NH_4_Cl and 0.017 M Tris-HCl (pH 7.2) for 1 min at room temperature and cell suspensions were washed and stained with relevant mAbs, as indicated below. Tumor tissue was mechanically disaggregated using a McIlwain tissue chopper (Mickle Laboratory Engineering), and a single-cell suspension was prepared by digesting the tissue in collagenase type A (Roche) and 25 μg/ml DNase (Sigma) in serum-free DMEM medium for 45 min at 37°C. Enzyme activity was neutralized by addition of DMEM containing 8% FCS, and the tissue was dispersed by passing through a 70-μm cell strainer. Lymphoid tissue was dispersed into single cells passing it through a 70-μm cell strainer. Single cells were first stained with APC-conjugated E7_59-57_/H-2D^b^ tetramers (Peptide and tetramer facility, Immunology department, Leiden University Medical Center) for 15 min at 4°C in the dark. After tetramer staining, tumor cells were incubated with 2% normal mouse serum with 10 µg/ml DNAse for 15 min on ice. For surface staining, cells were incubated for 30 min on ice in the dark with fluorochrome-conjugated antibodies and 0.5 *μ*l anti-APC mAb (clone APC003, BioLegend) per sample in PBS, 0.5% BSA, 0.01% sodium azide to increase intensity of tetramer staining. LIVE/DEAD^TM^ Fixable Near-IR Dead Cell Stain Kit (1:1000, Invitrogen) was added to exclude dead cells. Intracellular staining of cytokines and transcription factors was performed after cell fixation and permeabilization using the FOXP3 Transcription Factor Staining Buffer Set according to the manufacturer’s protocol (Thermo Fischer Scientific). The following fluorochrome-conjugated mAbs were used for flow cytometry and obtained from BD Pharmingen (Breda, the Netherlands) unless otherwise specified: CD45.2-BUV395 (1:200; clone 30-F11), TCRβ-PECy7 (1:100; clone H57-597), CD3-PECy7 (1:100, clone 145-2C11, eBiosciences), CD8-BUV805 (1:300, clone 53-6.7), CD4-BV711 (1:200, clone GK1.5), FOXP3-PECy5.5 (1:100, clone FJK-16s, eBiosciences), CD44-BV785 (1:200, clone IM7, BioLegend), CD62L-FITC (1:100, clone MEL-14, eBioscience), IFN*γ*-eF450 (1:200, clone XMG-1.2, eBioscience), Granzyme B-PE (1:200, clone GB11, Sanquin Amsterdam). To detect cytokine production, whole cell preparations from tumor and lymphoid tissue were plated in 100 µl IMDM/8% FCS in a 96-well U-bottom plate. Cells were treated with 50 ng/ml phorbol 12-myristate 13-acetate (PMA, Sigma Aldrich, Zwijndrecht, The Netherlands) and 1 µM ionomycin (Sigma Aldrich) dissolved in DMSO and diluted in culture medium. Control (unstimulated) cells were treated with an equal volume of DMSO diluted in culture medium. GolgiPlug (1 µg/ml; BD Biosciences) was added to all wells before incubating the cells for 3 hr at 37°C, 5% CO2. To determine absolute cell numbers AccuCount Blank Particles (Spherotech) were added to the samples, prior to analysis. For all experiments, cells were analyzed using a BD Symphony A5 flow cytometer with Diva software, and the generated data were analyzed using FlowJo software.

### Immuno-histochemical analysis

Harvested tumors were fixed for 24 hrs in aqueous solution with 50% ethanol, 5% acetic acid and 3.7% formalin, embedded in paraffin, and then sectioned randomly at 5 mm. For immunostaining, sections were rehydrated and incubated with primary antibodies to CD8 (eBioscience; clone 4SM15), CD4 (eBioscience; clone 4SM95) and FOXP3 (eBioscience; clone FJK-16s). Endogenous peroxidases were blocked with 3% H_2_O_2_, and the sections were then incubated with biotin-conjugated secondary antibodies, followed by incubation with HRP-conjugated streptavidin-biotin (DAKO). The substrate was developed using diaminobenzidine (DAB) (DAKO). We included negative controls to determine background staining, which was negligible. For the assessment of immune-cell infiltration in the tumor cross-sections, the immuno-stained slides were scanned and digitally processed using the Pannoramic 1000 scanner (3DHISTECH, Hungary) equipped with a 20x objective. Digital whole slide images of CD8-, CD4- and FOXP3-stained serial tissue sections were registered with the HE sections in HALO™ image analysis software version (3.2.1851.229) (Indica Labs, Corrales, NM). The tumor area within the stained sections were manually annotated and all nuclei within the tumor area (hematoxylin and/or DAB staining) were automatically segmented with the use of the commercially available Indica Labs – Multiplex IHC v2.3.4 algorithm module. Optimized parameters for the detection of nuclei signal included nuclear weight (1 for hematoxylin and 0.066 for DAB staining), nuclear contrast threshold (0.44), minimum nuclear optical density (0.095), nuclear size (11.3 – 220.7), minimal nuclear roundness (0.033) and nuclear segmentation aggressiveness (0.536). The optimized module parameters for the cytoplasmic and membrane detection included DAB-markup color (198, 163, 122) with the DAB-nucleus positive threshold (0.1105, 2.5, 2.5). The algorithm module parameters were kept constant for the analysis of all the sections across the different lymphocyte stainings. Next, with the utilization of the algorithm the total cell number within the tumor area (per section per staining) was automatically determined along with the equivalent number of each lymphocyte classification as DAB-positive cells. For the quantification analysis, the fraction (percentage) of DAB-positive cells (determined either via nucleus or cytoplasmic/membrane staining) over the total number of cells within the tumor area was used.

### Single-cell RNA-seq analysis of human melanoma tumors

Single-cell data from 32 melanoma tumors (Jerby-Arnon et al., 2018) was downloaded from NCBI GEO (gse115978) and exported to the R2 platform (http://r2.amc.nl, Mixed Melanoma SC – Regev - 7186 - tpm - gse115978). tSNE clustering was applied to 7186 cells. A complexity of 5 was chosen to represent the cohort. Inferred cell type information was extracted from the GEO dataset. Expression of ENPP2 and other annotations were projected onto the tSNE embedding. In every patient sample, the percentage of ENPP2-expressing cells was correlated to the percentage of cells inferred to be CD8^+^-positive. All analyses of the single-cell data were performed in the R2 genomics analysis and visualization platform.

### Statistical analyses

For *in vitro* migration assays, a two-tailed unpaired Student’s t-test was applied. A P value < 0.05 was considered statistically significant; *, p<0.05; **, p<0.005; ***, p<0.001; and ****, p<0.0001. Data from mouse studies were analyzed using GraphPad Prism version 9 (GraphPad Software, La Jolla, CA). Differences between various treatment groups were analyzed using the Mann-Whitney *U* Test. Differences in survival curves were analyzed using the Log Rank (Mantel-Cox) test. Differences with *P* values <0.05 were considered statistically significant.

## Authors’ Contributions

E.M.-R.: investigation, analysis, methodology, validation, visualization and writing; E.F.: investigation, methodology, analysis, validation, visualization and writing; I.v.d.H.A.: investigation, methodology, analysis, validation, and visualization; A.M.: investigation, methodology, data analysis and visualization; M.v.Z.: investigation, methodology, analysis, validation, and visualization; A.J.M.: investigation, methodology, and analysis; J.K.: investigation, methodology, analysis, and visualization; F.S.-P.: investigation, methodology, analysis, and visualization; S.K.: methodology and analysis; T.L.: methodology and analysis; J.H.: conceptualization and methodology; Z.J.: conceptualization, analysis, and methodology; T.N.S.: conceptualization, methodology; A.P.: analysis, methodology, and validation; I.V.: conceptualization, investigation, analysis, methodology, validation, visualization, and writing; J.v.d.B.: conceptualization, investigation, analysis, methodology, validation; J.B.: conceptualization, analysis, methodology, supervision and writing; W.H.M.: conceptualization, analysis, methodology, validation, supervision, and writing.

## Acknowledgments

We thank Paula Voabil, Jos Urbanus, Yanling Xao, Marjolein Mertz and Juan Manuel Alba for their help with experiments and data analysis, Kees Franken for making MHC tetramers, Tomasz Ahrends and Anne van der Leun for helpful discussions and sharing unpublished data. This work was supported by private funding to W.H.M and grants from the Dutch Cancer Society (grants NKI 2013-5951 and 10764 to I.V.; grant NKI 2017-10894 to J.B. and I.V.); the German research foundation (DFG; grant ME 4924/1-1 to A.M; and the NIH (grant P30 GM127211 to A.J.M). E.M.R is supported by an ‘Ramón y Cajal’ Award (RYC2019-027950-I) from Ministerio de Ciencia e Innovación (MINECO).

---

**Figure S1.**
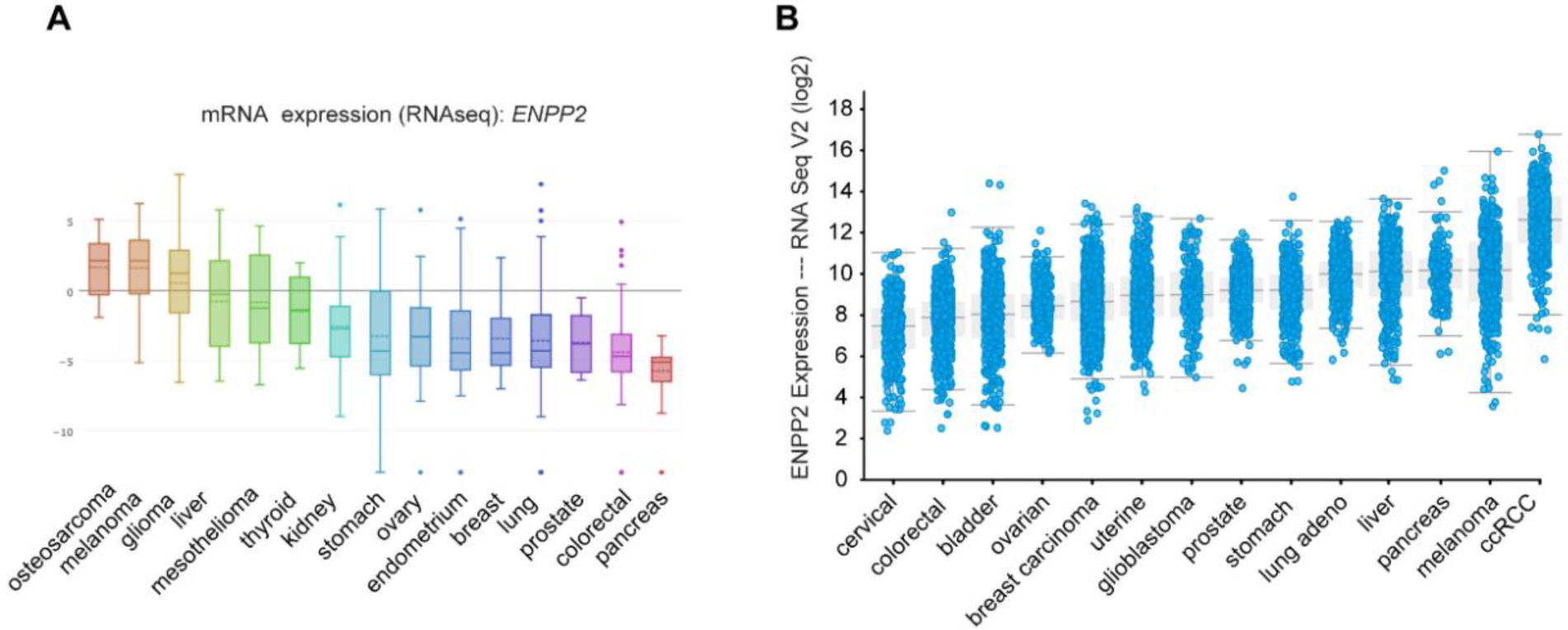
ATX mRNA expression in cancer cell lines and solid tumors. ***(A****) ENPP2* expression in the indicated cell lines ranked according to median values. Note high *ENPP2* expression in melanoma cell lines (n=61). RNAseq expression data were retrieved from the Cancer Cell Line Encyclopedia (CCLE; https://portals.broadinstitute.org/ccle). **(B)** Pan-cancer analysis of *ENPP2* expression in the indicated solid tumors ranked according to median values (ccRCC, clear cell renal cell carcinoma). RNAseq v2 mRNA expression data were retrieved from the TCGA database (www.cbioportal.org). Note that *ENNP2* expression in cancer cell lines poorly correlates with that in the corresponding tumors, which is attributed to the presence of ATX-expressing stromal cells.

**Figure S2.**
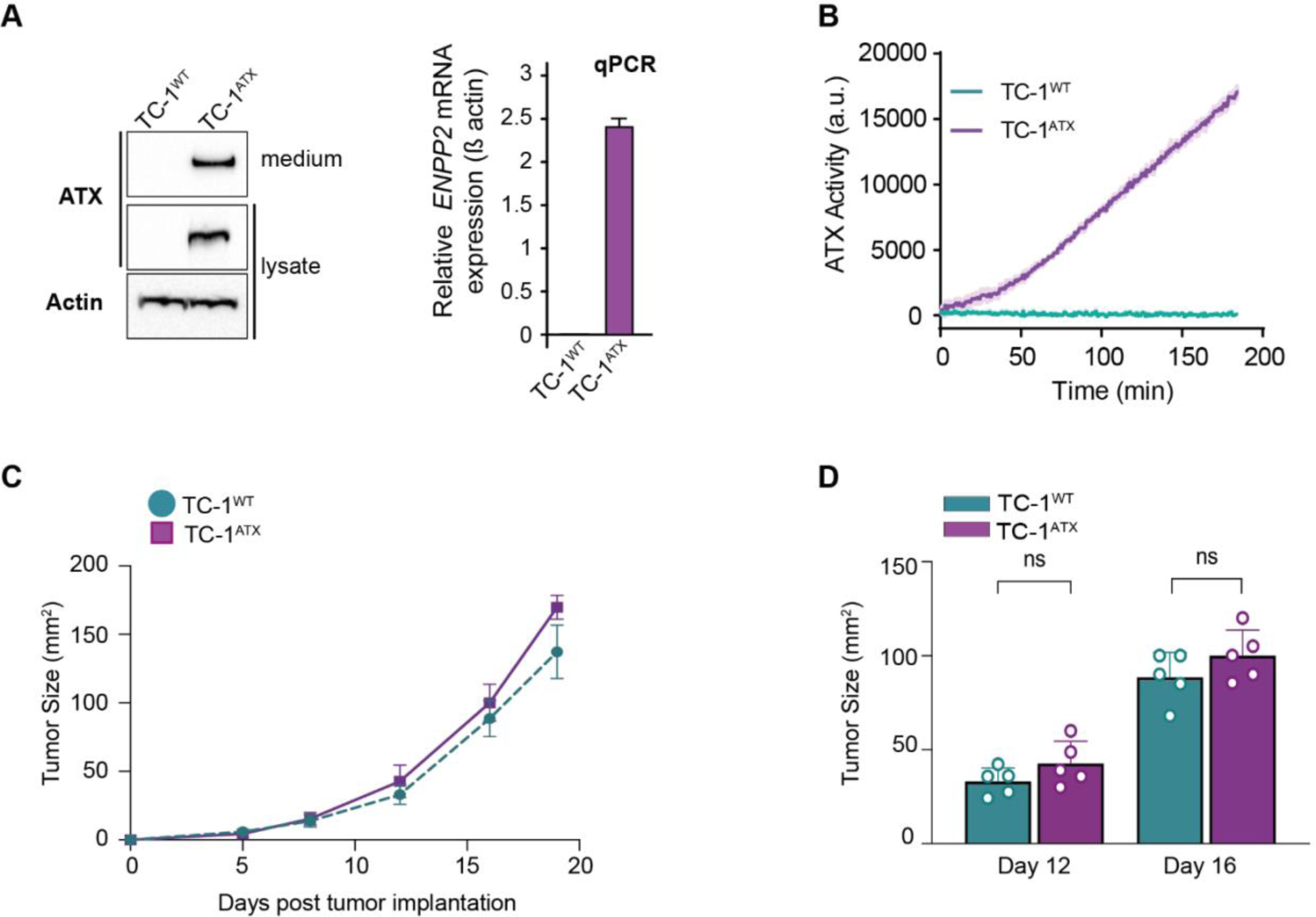
Characteristics of wild-type and ATX-expressing TC-1 tumors. **(A)** (<u>Left</u>) immunoblot analysis of ATX protein expression in wild-type (TC-1^WT^) and ATX-expressing (TC-1^ATX^) tumor cells. Actin was used as loading control. (<u>Right</u>) ATX mRNA expression (relative to Cyclophilin) in TC-1^WT^ and TC-1^ATX^ cells as analyzed by qPCR. **(B)** Secreted ATX (lysoPLD) activity in supernatants from TC-1^WT^ and TC-1^ATX^ cells, as measured by choline release from added LPC(18:1) over time. See Methods for details. **(C)** TC-1^WT^ and TC-1^ATX^ tumor growth expressed as mean size in non-vaccinated mice (n=5). **(D)** Average tumor size in the same mice as in (**C**) on days 12 and 16 after s.c. tumor cell implantation. Data is depicted as mean ± SD; ns: not significant (Mann-Whitney U test).

**Figure S3.**
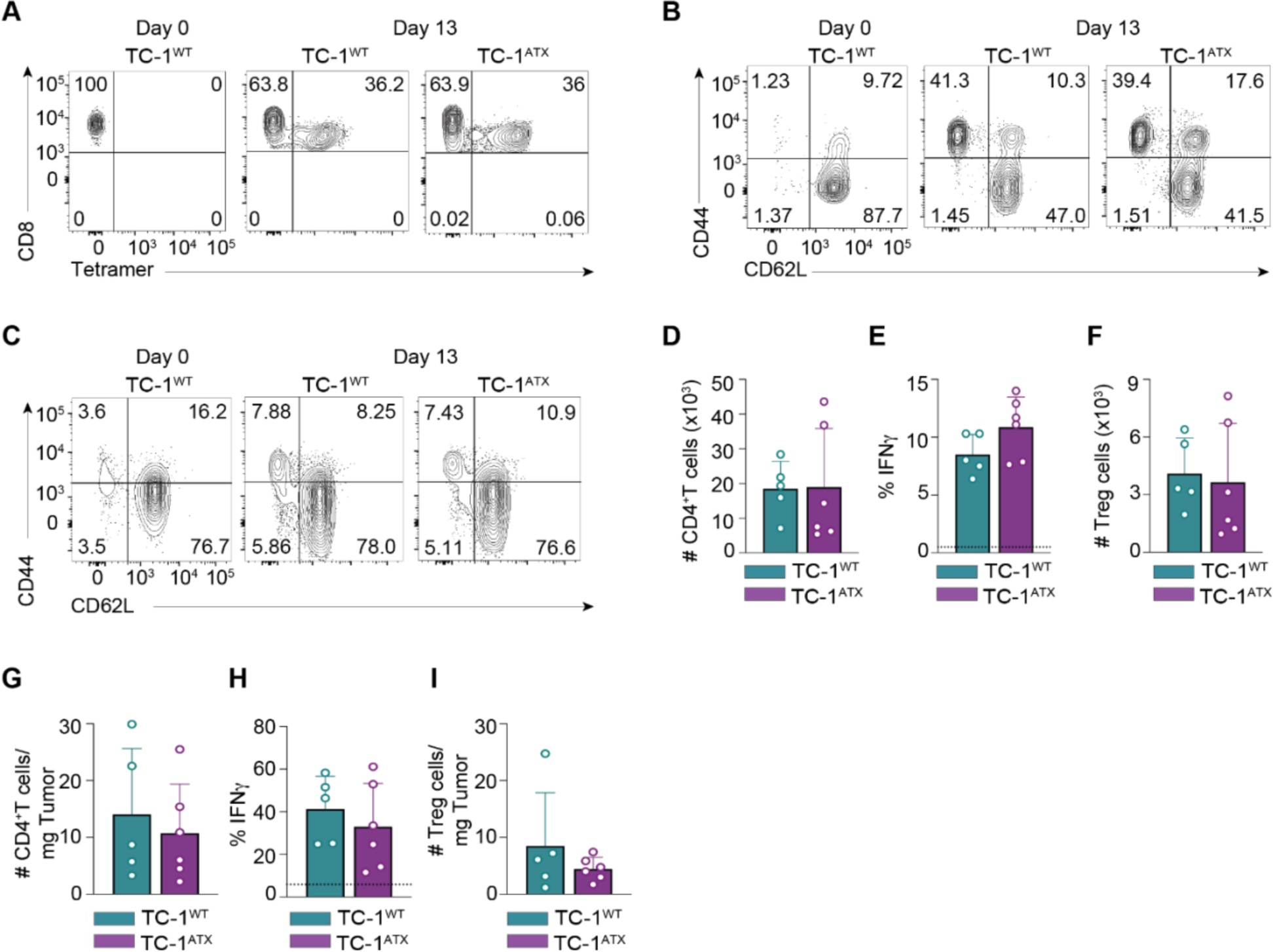
Enforced ATX expression in tumor cells does not affect the CD4^+^ T cell response to vaccination. **(A-C)** Primary data belonging to Figure 5. Representative flow cytometry plots depicting H-2D^b^/E7_49–57_ Tet^+^ cells **(A)** and CD44^+^CD62L^-^ effector phenotype cells among total CD8^+^ T cells **(B)** and total CD4^+^ T cells **(C)** in blood from TC-1^WT^ (n = 6) and TC-1^ATX^ (n = 5) tumor-bearing mice at day 13 post vaccination. Data at day 0 are from non-vaccinated TC-1^WT^ tumor-bearing mice. **(D-I)** CD4^+^ T cell populations as analyzed by flow cytometry in spleen (**D-F**) and tumors (**G-I**) of TC-1^WT^ (n = 5) and TC-1^ATX^ (n = 6) tumor-bearing mice at day 18 after tumor implantation. **(D)** Absolute number (#) of FOXP3^-^ CD4^+^ conventional T cells (Tconv) in spleen. **(E)** Frequency of IFNᵧ^+^ cells among conventional CD4^+^ T cells in spleen. IFNᵧ was measured as outlined in Figure 5. **(F)** Absolute number (#) of FOXP3^+^ CD4^+^ T cells (Tregs) in spleen. **(G)** Absolute number (#) of CD4^+^ Tconv cells per mg tumor tissue, found in TC-1^WT^ and TC-1^ATX^ tumor-bearing mice. **(H)** Frequency of IFNᵧ^+^ cells within CD4^+^ Tconv cells in the tumors. IFNᵧ was measured as outlined in Figure 5. **(I)** Absolute number (#) of CD4^+^ Tregs in TC-1^WT^ and TC-1^ATX^ tumors. **(D-I)** Data are depicted as mean ± SD; no significance for all cell populations analyzed (Mann-Whitney U test). Data are from one experiment representative of two experiments.

**Figure S4.**
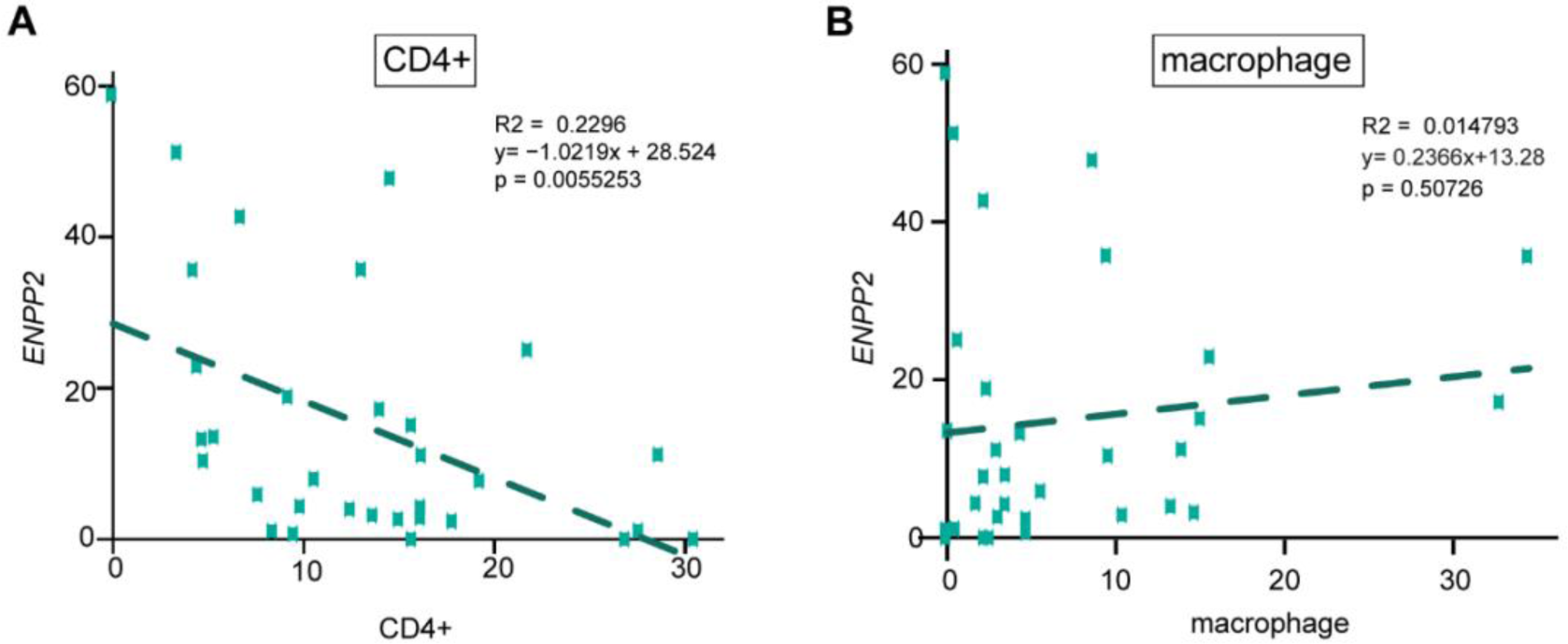
*ENPP2* expression and accumulation of CD4^+^ T cells (A) and macrophages (B) in melanoma tumors (n=32). Pearson’s correlation between the percentage of inferred *ENPP2*-expressing cells and CD4^+^ T cells versus macrophages as indicated. For details see text and Figure 7.

